# Neural Correlates of Social Withdrawal and Preference for Solitude in Adolescence

**DOI:** 10.1101/2025.03.17.643680

**Authors:** Matthew Risner, Catherine Stamouls

## Abstract

Social isolation during development, especially in adolescence, has detrimental but incompletely understood effects on the brain. This study investigated the neural correlates of preference for solitude and social withdrawal in a sample of 2,809 youth (median (IQR) age = 12.0 (1.1) years, 1440 (51.26%) females) from the Adolescent Brain Cognitive Development study. Older youth whose parents had mental health issues more frequently preferred solitude and/or were socially withdrawn (β=0.04–0.14, CI=[0.002,0.19], p<0.05), both of which were associated with internalizing and externalizing behaviors, depression, and anxiety (β=0.25-0.45, CI=[0.20,0.49], p<0.05). Youth who preferred solitude and/or were socially withdrawn had lower cortical thickness in network regions supporting social function (cuneus, insula, anterior cingulate and superior temporal gyri) and/or mental health (β=-0.09 to -0.02, CI=[-0.14,-0.003], p<0.05), and higher amygdala, entorhinal cortex, parahippocampal gyrus, and basal ganglia volume (β=2.62-668.10, CI=[0.13,668.10], p<0.05). Youth preferring solitude had more topologically segregated dorsal attention, temporoparietal, and social networks (β=0.07-0.10, CI=[0.02,0.14], p≤0.03). Socially withdrawn youth had less topologically robust and efficient (β=-0.05 to -0.80, CI=[-1.34,-0.01], p<0.03), and more fragile cerebellum (β=0.04, CI=[0.01,0.07], p<0.05). These findings suggest that social isolation in adolescence may be a risk factor for widespread alterations in brain regions supporting social function and mental health.

## 1. INTRODUCTION

Adolescence is a period of extensive physical, cognitive, and social changes, but also heightened neural maturation and circuit reorganization (Best and Ban, 2021; Blakemore and Mills, 2014a; Klimstra et al., 2010; Luna, 2009; Özdemir et al., 2016; Pfeifer and Allen, 2021; Rogol et al., 2002; Siervogel et al., 2004; Steinberg and Morris, 2001; Susman and Rogol, 2004; Wigfield et al., 2006; Andrews et al., 2021; Ashtari et al., 2007; Blakemore, 2008, 2012; Goddings et al., 2014; Lenroot et al., 2007; Mills et al., 2016; Peper et al., 2011; Sturman and Moghaddam, 2011; Vijayakumar et al., 2018; Wierenga et al., 2014; Yurgelun-Todd, 2007). Concurrently, the youth social world expands and changes, in part due to social reorientation –a shift in the relative importance of peer relationships compared to those with family (Albert et al., 2013; Giordano, 2003; Jaccard et al., 2005; Lerner and Steinberg, 2009; Maxwell, 2002; Sawyer et al., 2018; Wang et al., 1995). Changes in the youth social environment can be overwhelming, sometimes leading to social isolation (Barzeva et al., 2019; Biggs et al., 2012; Oh et al., 2008; Porcelli et al., 2019; Rubin et al., 2009; Wood et al., 2022).

Although spending time alone may be motivated by a desire for self-reflection or creative endeavors, habitual withdrawal driven by social stressors, for example persistent peer rejection (Borg and Willoughby, 2022; Vasey, 2001), can lead to social isolation, which can have profound adverse effects on social development (Almeida et al., 2021; Orben et al., 2020; Thompson et al., 2022). The adolescent brain is vulnerable to social stressors, which increase the risk of circuit miswiring, and mental health and behavioral issues (Blakemore, 2019; Bor et al., 2014; Costello et al., 2011; Reiss, 2013).

Negative factors in the youth’s immediate environment increase the risk for social withdrawal, isolation, and loneliness. Children whose parents have mental health issues may be at higher risk of behavioral and mental health problems, including social isolation (Manning and Gregoire, 2009; Matthews et al., 2015; Van Loon et al., 2014).

Stigma surrounding parental mental illness can cause children to face bullying, embarrassment, and guilt. This often leads to concealing their family circumstances and avoiding support, further exacerbating isolation (Bosch et al., 2017; Cogan et al., 2005; Dam et al., 2018; Reupert et al., 2021). Relationships with peers are similarly influential, and peer rejection, victimization, lacking friends, and low friendship quality also contribute to loneliness (Asher and Paquette, 2003; Cassidy and Asher, 1992; Cheng and Furnham, 2002; Lodder et al., 2017; Schwartz-Mette et al., 2020; Vanhalst et al., 2014; Woodhouse et al., 2012). These relationships are, however, bidirectional since social withdrawal may worsen the quality and/or depth of friendships (Barzeva et al., 2022; Biggs et al., 2012; Coplan et al., 2018; Rubin et al., 2006). Social exclusion at school may lead to social withdrawal and loneliness (G. Arslan et al., 2023; Ph. D. Arslan Gökmen, 2021; Ford et al., 2018; Patton et al., 2006). In contrast, a sense of school connectedness may mitigate the adverse psychological effects of social isolation (Foster et al., 2017; Hall-Lande et al., 2007; London and Ingram, 2018; Marraccini and Brier, 2017; Preston and Rew, 2022). These associations are bidirectional, as socially withdrawn children often encounter difficulties in school, including poor relationships with teachers, academic struggles, and may avoid school altogether. These challenges further reinforce their social isolation and limit opportunities for meaningful social engagement (Coplan et al., 2018; Rubin et al., 2009; Stenseng et al., 2022).

Lack of social interaction increases the risk of physical, cognitive, and mental health problems (Christiansen et al., 2021; Fuhrmann et al., 2019; Hall-Lande et al., 2007; Houghton et al., 2022; Rubin et al., 2009; Stenseng et al., 2022). Social isolation and loneliness have been associated with increased risk for cancer, obesity, diabetes, asthma, migraine, hypertension, and back pain in adolescents and young adults (Almeida et al., 2021; Christiansen et al., 2021; Goosby et al., 2013; Mahon et al., 1993; Mushtaq et al., 2014; von Soest et al., 2020). They have also been associated with longer-term physical issues, including obesity, cardiovascular problems, inflammation, and impaired immunoregulation in adulthood (Caspi et al., 2006; Danese et al., 2009; Goosby et al., 2013; Hawkley et al., 2010; Hawkley and Cacioppo, 2010; Lacey et al., 2014; Matthews et al., 2024; Patterson and Veenstra, 2010).

Social isolation may also significantly increase the risk of mental health issues, In adolescence, during which many common mental health disorders emerge (Blakemore, 2019; de Girolamo et al., 2012; Paus et al., 2008), it has been linked to anxiety, depression, internalizing problems, and suicidal ideation and self-harm (Blakemore, 2019; Bor et al., 2014; Cacioppo et al., 2006; Costello et al., 2011; Endo et al., 2017; Gallagher et al., 2014; Harman et al., 2021; Jones et al., 2011; McClelland et al., 2020; Reiss, 2013; Rubin et al., 2009; Schinka et al., 2012). Social isolation has also been associated with cognitive deficits, including worse academic performance and impaired reward processing and language skills (Almeida et al., 2021; Jefferson et al., 2023; Matthews et al., 2023; Rubin et al., 2009; Tomova et al., 2022). In addition, recent studies focusing on social isolation of youth during the COVID-19 pandemic have identified associations between pandemic-related social isolation and deficits in executive function, attention, and memory (Lavigne-Cerván et al., 2021; Murtaza et al., 2023).

The neural correlates of social isolation (resulting from social withdrawal and/or preference for solitude) in the adolescent brain remain poorly understood. This is a significant gap in knowledge given that adolescence represents a formative period for the establishment of social identity and mental health outcomes (Blakemore, 2019; Blakemore and Mills, 2014b; Branje et al., 2021; Fergusson et al., 2005; Kessler et al., 2005; Kroger, 2004; Meeus, 1996; Paus et al., 2008; Schlack et al., 2021). Adult studies have shown that loneliness and social withdrawal are associated with lower gray matter volume in the left posterior superior temporal, the amygdala, hippocampus and cerebellum (Düzel et al., 2019; Kanai et al., 2012; Koolschijn et al., 2013; Lind et al., 2020), disrupted white matter integrity in the salience network (Tian et al., 2014), and lower white matter density in the inferior parietal lobule and anterior insula (Lam et al., 2021; Nakagawa et al., 2015). It has also been associated with changes in gray matter volume and lower white matter density in the dorsolateral prefrontal cortex, likely leading to emotional dysregulation (Kong et al., 2015; Liu et al., 2016; Zheng et al., 2023). Social isolation may also affect emotional processing in adolescents through its impact on anatomical connections between the amygdala and the orbitofrontal cortex (Goetschius et al., 2020). Together, these findings suggest that social isolation adversely impacts the structural integrity of brain regions that play critical roles in social perception, cognitive control, and emotional processing.

Beyond its impact on brain structure, social withdrawal, isolation and loneliness in young adults has been linked to higher resting-state functional connectivity in the right central operculum, supramarginal gyrus, and circuits involved in sustained attention (Layden et al., 2017), and weaker connections between dorsal attention and salience networks, as well within and between temporal, limbic, and prefrontal networks (Feng et al., 2019; McIver et al., 2019; Tian et al., 2017). In adolescents, social withdrawal has been associated with aberrant stronger connections between the left amygdala and regions of the dorsal attention, frontoparietal control, and auditory networks, and between the prefrontal and limbic networks (Thomas et al., 2024). Loneliness been associated with lower activation of the ventromedial prefrontal cortex (Golde et al., 2019) and higher functional connectivity in visual attention networks (Brilliant T. et al., 2022).

Despite valuable insights provided by prior studies, adolescent brain correlates of social isolation (as a result of withdrawal and/or preference for solitude) remain incompletely understood and have not been systematically characterized. A significant barrier to robust investigations is the heterogeneity of adolescent brain development, and the complexity of associations between social behaviors, environmental factors, and the brain. However, the availability of multimodal (including neuroimaging) data from the Adolescent Brain Cognitive Development (ABCD) study, a historically large (∼12,000 youth) investigation of adolescent brain development (Casey et al., 2018) provides a unique opportunity to bridge this gap in knowledge, and investigate relationships between social isolation and brain development during this complex period.

The present study leveraged the ABCD structural MRI and resting-state fMRI data, mental health assessments and survey data on social behaviors and environmental factors, in order to investigate associations between preference for solitude and social withdrawal (both leading to social isolation), and the organization of intrinsically coordinated brain networks, and their structural correlates in almost 3,000 adolescents (ages 11-12 years), who at this age may be at increased risk of mental health disorders and maladaptive social behaviors (Kessler et al., 2005). The study hypothesized that preference for solitude and social withdrawal are in part driven by negative factors in the youth social environment and are associated with widespread structural and functional brain changes, especially in underdeveloped regions involved in social function.

## 2. MATERIALS AND METHODS

This study analyzed publicly available and thus anonymized data, and was approved by the Institutional Review Board.

### 2.1 Participants

A sample of n = 2809 youth from the ABCD study, measured at the two-year follow-up, was analyzed. The primary criterion for inclusion was availability of at least one high-quality (≤10% of frames censored for motion) 5-min resting-state fMRI run that had been collected prior to the onset of the COVID-19 pandemic (assumed to be March 11, 2020, when the World Health Organization declared it as such). Youth measured during the pandemic were excluded to eliminate its potential confounding effects on the brain and social behaviors. Although studying the effects of social isolation during the pandemic on the brain is critical, it was outside the scope of the present study. In addition, participants with clinical MRI findings and neuropsychiatric or neurodevelopmental disorders were also excluded, since they may independently affect brain morphology and the organization of neural circuits (Brooks et al., 2021, 2022, 2023). The final analytic sample included 1366 biological boys (48.63%) and 1440 girls (51.26%), with a median age of 12.0 years (Interquartile Range (IQR) = 1.1 years). Over half of youth were white non-Hispanic (1488 (53.0%)), 1069 (38.1%) were non-Hispanic racial minorities, and 243 (8.7%) were Hispanic. Socioeconomic status was assessed based on annual household income, with a median income $75,000-$99,999. The sample included 515 (18.3%) youth in pre-puberty, 609 (21.7%) in early puberty, 918 (32.7%) in mid-puberty, and 622 (22.1%) in late/post-puberty. Detailed demographic and other participant characteristics are provided in Table 1.

**Table 1.**
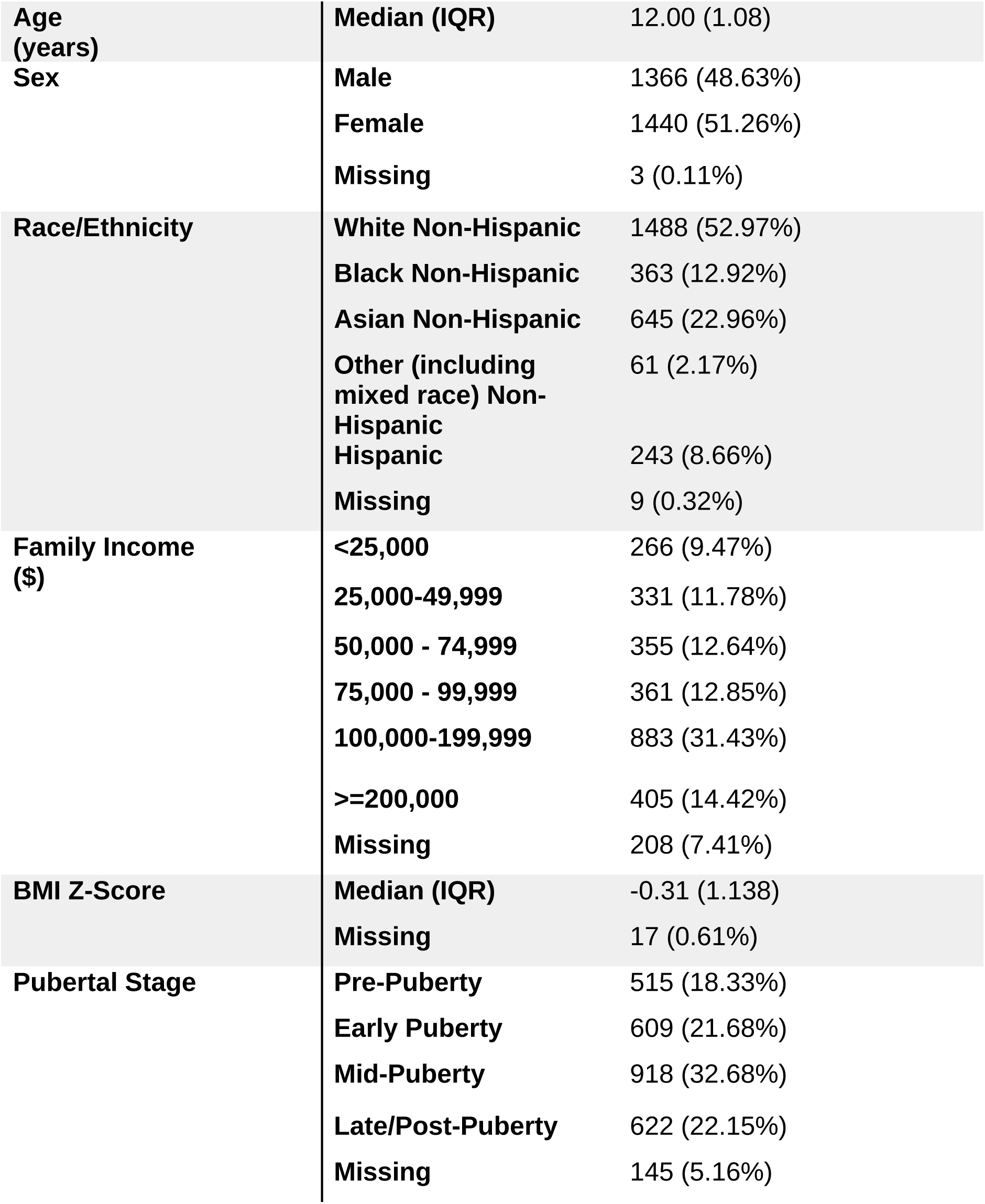
Demographic and other individual characteristics (n = 2809).

|  |  |  |
| --- | --- | --- |
| **Age (years)** | **Median (IQR)** | 12.00 (1.08) |
| **Sex** | **Male** | 1366 (48.63%) |
|  | **Female** | 1440 (51.26%) |
|  | **Missing** | 3 (0.11%) |
| **Race/Ethnicity** | **White Non-Hispanic** | 1488 (52.97%) |
|  | **Black Non-Hispanic** | 363 (12.92%) |
|  | **Asian Non-Hispanic** | 645 (22.96%) |
|  | **Other (including mixed race) Non-Hispanic** | 61 (2.17%) |
|  | **Hispanic** | 243 (8.66%) |
|  | **Missing** | 9 (0.32%) |
| **Family Income (\$)** | **<25,000** | 266 (9.47%) |
|  | **25,000-49,999** | 331 (11.78%) |
|  | **50,000 - 74,999** | 355 (12.64%) |
|  | **75,000 - 99,999** | 361 (12.85%) |
|  | **100,000-199,999** | 883 (31.43%) |
|  | **>=200,000** | 405 (14.42%) |
|  | **Missing** | 208 (7.41%) |
| **BMI Z-Score** | **Median (IQR)** | -0.31 (1.138) |
|  | **Missing** | 17 (0.61%) |
| **Pubertal Stage** | **Pre-Puberty** | 515 (18.33%) |
|  | **Early Puberty** | 609 (21.68%) |
|  | **Mid-Puberty** | 918 (32.68%) |
|  | **Late/Post-Puberty** | 622 (22.15%) |
|  | **Missing** | 145 (5.16%) |

### 2.2 Measures of social withdrawal

Two measures directly assessing social withdrawal and preference for solitude were extracted from the Child Behavior Checklist (CBCL), completed by parents: *“withdrawn, doesn’t get involved with others”* and *“would rather be alone than with others.”* Both were rated on a Likert scale from not true (0) to very/often true (2).

### 2.3 Measures of mental health

Data on common mental health problems that were adequately represented in the study sample, including anxiety, depression, internalizing and externalizing behaviors (all t-scores) were extracted from the CBCL. Additional information on social anxiety disorder and anhedonia was extracted from the Kiddie Schedule for Affective Disorders and Schizophrenia (K-SADS), also completed by parents. For each mental health factor from the K-SADS, responses to two yes/no questions about past and present diagnoses or symptoms were combined into a single binary variable based on any positive responses.

### 2.4 Environmental factors

Multiple aspects of the youth social environment were investigated. Parental mental health was based on the Adult Behavior Checklist (ABCL), and was measured as a t-score of critical items related to depression, self-harm, substance use, mood swings, and suicidal thoughts. Family dynamics were assessed based on the parent-reported Family Environment Scale. Conflict and cohesion were measured on continuous scales by aggregating responses from multiple survey items designed to assess specific aspects of the family’s organizational structure. In addition, parent responses to one question from the Mexican American Cultural Values Scale were extracted, providing a measure for how strongly they believed that family members should show love and affection to one another. Responses were coded on a Likert scale from not at all (1) to completely (5). Prior work has identified associations between this parental belief youth prosocial behaviors (Smith and Stamoulis, 2023).

Questions on peer relations were: 1) number of close friends (from the Other Resilience survey); 2) regular group of friends (binary variable, from the KSADS); 3) bullying, represented as a binary variable based on questions on whether a child had been bullied in any social setting (from a parent-reported K-SADS question) or had been cyberbullied (from the youth-reported Cyber-bullying instrument). The Discrimination Measure, reported by youth, assessed experiences of perceived discrimination in four domains: race/ethnicity/color, being from another country, sexual orientation, and/or weight. Given small samples with positive responses in each domain, responses were combined into a single binary variable for analysis, indicating any type of discrimination. Youth responses on whether they got along with their teacher and liked school (from the School Risk and Protective Factors survey) were also examined. Responses were coded on a Likert scale from definitely not true (1) to definitely true (4).

### 2.5 Additional characteristics

Other assessments of youth social behaviors were extracted from The Early Adolescent Temperament Questionnaire (completed by the parents) and included *“Is energized by being in large crowds of people,” “Wants to have close relationships with other people,”*

*“Is shy,”* and *“Feels shy about meeting new people”*, all on a 5-point scale (almost always untrue (1) to almost always true (5)). Another question from the same instrument asked if the child *“Likes meeting new people”*, and was reverse-coded by the ABCD, with higher values indicating less frequent social engagement. Two questions on past or present symptoms of a fear of social situations were extracted from the KSAD-S and combined into a binary variable with 1 corresponding to a ‘true’ response to either or both questions, and 0 otherwise. Another question from the CBCL asked parents if their child was *“self-conscious or easily embarrassed”* and was coded on a Likert scale from not true (0) to very/often true (2).

### 2.6 Neuroimaging data analysis

#### 2.6.1. Resting-state fMRI processing and topological property estimation

Individual structural MRI and resting-state fMRI (rs-fMRI; participants completed up to four 5-minute runs), underwent initial preprocessing by the ABCD study’s Data Analysis, Informatics and Resource Center (DAIRC), to correct for bias field, B0 distortion, grad warp and motion in the scanner (Hagler et al., 2019)). Additional processing was necessary to further suppress motion-related and other artifacts, harmonize the data across scanners (data were collected with a 3T GE, Siemens or Philips scanner), and downsample the voxel-level time series. The custom Next Generation Neural Data Analysis (NGNDA) pipeline was used for this purpose (Next-Generation Neural Data Analysis (NGNDA), 2020). Following segmentation of each participant’s structural MRI (sMRI), coregistration of their fMRI to the sMRI, and normalization to common MNI152 space, fMRI signals were further processed to improve their quality. This process included removal of initial frames, correction for motion, and denoising using signal decomposition to eliminate signal components likely associated with cardiorespiratory and other artifacts (Brooks et al., 2021). Voxel-level time series were then downsampled using three atlases for parcellating cortical and subcortical structures and the cerebellum, resulting in 1088 parcel signals (Diedrichsen et al., 2009; Schaefer et al., 2018; Tian et al., 2020). Only fMRI runs with ≤10% motion-censored frames were retained for further analysis. Each participant’s best-quality run (based on lowest median connectivity, given that the brain at rest is overall weakly coordinated at rest) was first analyzed. Often this run also had the lowest number of frames censored for motion. For replication purposes, the second best-quality run was selected from a subsample of participants (n = 2160 (∼77%) of the sample) who had more than one run of adequate quality for analysis. Median (IQR) percent of frames censored for motion was 1.07% (IQR = 4.00%) in the best-quality run and 1.07% (IQR = 3.73%) in the second run. Associations between resting-state topological properties and measures of social withdrawal that were consistent across runs are reported in primary analyses and tables. Additional associations based on the best-quality run, and also larger sample, are reported in secondary analyses and supplemental tables.

Resting-state connectivity was calculated as the peak cross-correlation between pairs of fMRI signals, resulting in symmetric 1088-by-1088 matrices. All methods used to estimate connectivity yield full matrices that include spurious correlations that need to be eliminated prior to the estimation of topological properties. In this study, statistical thresholds were estimated from the entire cohort and all available runs, and bootstrapping with replacement was used to derive thresholds for different levels of connection strength. A conservative threshold, equal to the moderate outlying value (median + 1.5*IQR) of peak cross-correlation was then used to obtain adjacency matrices. Topological properties were estimated from these matrices at multiple spatial scales: a) brain regions (network nodes), b) large-scale networks that included the reward and social networks (Yeo et al, 2011; Haber and Knudson, 2010, Blakemore, 2008), c) entire brain (the connectome). Topological properties at the network and connectome levels included modularity, global clustering, global efficiency, natural connectivity (robustness), median connectivity (within and between networks), fragility, stability, and segregation. The latter was calculated as the ratio of within-network over out-of-network median connectivity. Node properties included degree, local clustering, and eigenvector centrality. Detailed descriptions of each of these properties are provided elsewhere (Pasqualetti et al., 2020; Rubinov and Sporns, 2010; Wu et al., 2011).

#### 2.6.2. Structural parameters

Morphometric brain properties (provided by the ABCD) were analyzed. These were estimated using a parcellation based on the Desikan-Killiany atlas, which segments the cortex into 68 regions based on anatomical landmarks (Desikan et al., 2006).

Subcortical voxels were parcellated into 30 regions using a probabilistic method (Fischl et al., 2002). Properties of 98 regions were analyzed, including cortical thickness and white matter intensity, and cortical and subcortical volume (Hagler et al., 2019). White matter intensity was was also examined as a potentially sensitive cortical marker of brain development (Hagler et al., 2019; Lewis et al., 2018; Westlye et al., 2009).

### 2.7 Statistical analysis

All relationships of interest were investigated using linear mixed-effects regression models that included a random intercept and slope for each of the 21 ABCD sites to account for potential site effects. All models included age, pubertal stage, sex, race-ethnicity, family income, and BMI z-score stratified by sex (which has been previously linked to differences in topological properties in the ABCD cohort (Brooks et al., 2023). All analyses accounted for potential bias associated with sampling differences at the 21 sites, using the ABCD-provided propensity scores. Given limited statistical power to compare individual race/ethnic groups, race-ethnicity was coded as a binary variable representing white non-Hispanic participants (0) and racioethnic minorities (1). Pubertal stage was coded as an ordinal variable in the range 1 (pre-puberty) to 4 (late/post-puberty).

In models that included brain parameters (the dependent variables), social withdrawal and preference for solitude were the independent variables of interest. These models were also adjusted for internalizing behavior scores, a variable that was consistently significant across analyses. In models that examined association between social isolation and environmental, mental health, and other behavioral factors, social withdrawal and preference for solitude were the dependent variables. Finally, models with resting-state network parameters included percent of frames censored for motion and scan time of day participants (0-23 h) as additional variables (Brooks et al., 2021; Hu et al., 2023; Vaisvilaite et al., 2022). Variations of models were also developed, in order to assess pairwise comparisons between specific response categories of the social withdrawal measures, focusing on two categories at a time, or treating these measures as binary variables.

The significance level was set at ɑ = 0.05. All parameter p-values were adjusted for the False Discovery Rate (FDR), using a well-established method (Benjamini and Hochberg, 1995). In analyses focusing on topological properties of the entire brain or individual networks, the FDR correction was done over properties. In analyses focusing on regional (node) properties, the correction was done over properties of all nodes within a particular network. P-values for parameters in structural models were adjusted over regional morphometric properties. Associations were considered significant if the model intercept, parameter of interest, and model p-value were all significant. Model cross-validation was performed by randomly dividing the data into training and testing sets (75:25) 100 times. The coefficient of variation of the root-mean-squared error (CV[RMSE]) between predicted and observed responses was used to assess each model’s predictive power. All data were analyzed in the Harvard Medical School high-performance cluster using the software MATLAB (release R2023a, Mathworks, Inc).

## 3. RESULTS

Across the cohort, 455 (17.9%) youth sometimes or often preferred solitude, and 170 (6.0%) were sometimes or often socially withdrawn. Less than 10% reported past or present anhedonia (264 (9.4%)) and 40 (1.4%) were diagnosed with social anxiety disorder. Median (IQR) t-scores for other mental health measures were 51.0 (2.0) for anxiety, 50.0 (4.0) for depression, 41.0 (16.0) for externalizing behaviors, and 46.0 (14.0) for internalizing behaviors.

Median (IQR) parent mental health score was 51.0 (4.0). Almost 90% of parents said that it was important for the family to show love and affection towards each other. Median family cohesion was 8.0 (2.0) whereas median family conflict was 2.0 (2.0). Median (IQR) family size was 4 (1). Participants had on average 5 close friends, and almost 90% had a regular group of kids to hang out with. About 10% reported being discriminated against (302 (10.7%)), and 519 (18.5%) had been bullied. Almost 95% reported having good relationships with teachers, and ∼70% said that they liked school a lot (1991 (70.9%)). Summary statistics for social environmental factors are provided in Table 2.

**Table 2.** Distribution of social environmental factors in the cohort.

| **PARENTAL FACTORS** |  |  |
| --- | --- | --- |
| Parental mental health (T-score) | Median (IQR) | 51 (4) |
|  | Missing | 378 (13.46%) |
| Parental belief that family members should show love and affection to one another | Not at all | 15 (0.53%) |
|  | A little | 23 (0.82%) |
|  | Somewhat | 254 (9.04%) |
|  | Very much | 1237 (44.04%) |
|  | Completely | 1279 (45.53%) |
|  | Missing | 1 (0.04%) |
| **FAMILY ENVIRONMENT** |  |  |
| Family conflict Scale: 0-9 | Median (IQR) | 2 (2) |
|  | Missing | 1 (0.04%) |
| Family cohesion Scale: 0-9 | Median (IQR) | 8 (2) |
|  | Missing | 1 (0.04%) |
| Family size | Median (IQR) | 4 (1) |
|  | Missing | 36 (1.28%) |
| **PEER RELATIONSHIPS** |  |  |
| Number of close friends | Median (IQR) | 5 (5) |
|  | Missing | 6 (0.21%) |
| Has a regular group of kids to hang out with at school/neighborhood | No | 276 (9.83%) |
|  | Yes | 2427 (86.40%) |
|  | Missing | 106 (3.77%) |
| Faced discrimination in the past 12 months | No | 2507 (89.25%) |
|  | Yes | 302 (10.75%) |
|  | Missing | 0 (0%) |
| Problems with bullying at school/neighborhood OR cyberbullying | No | 2276 (81.03%) |
|  | Yes | 519 (18.48%) |
|  | Missing | 14 (0.50%) |
| **SCHOOL** |  |  |
| Gets along with their teachers | Strongly disagree | 25 (0.89%) |
|  | Disagree | 120 (4.27%) |
|  | Agree | 1188 (42.29%) |
|  | Strongly agree | 1471 (52.37%) |
|  | Missing | 5 (0.18%) |
| Likes school a lot | Strongly disagree | 259 (9.22%) |
|  | Disagree | 554 (19.72%) |
|  | Agree | 1204 (42.86%) |
|  | Strongly agree | 787 (28.02%) |
|  | Missing | 5 (0.18%) |

Almost 70% of parents reported that their child wanted close relationships with other people (1938 (69.0%)), ∼60% of youth were not self-conscious or easily embarrassed (1717 (61.%), and liked to meet new people (1654 (58.9%)), and almost 30% were energized by large crowds (813 (28.9%)). About a quarter were shy (552 (26.4%)), fewer were specifically shy about meeting new people (466 (16.6%)), and even fewer were afraid of social situations (150 (5.3%)). Detailed distributions of responses are provided in Table S1.

### 3.1 Associations between social isolation, behaviors and mental health outcomes

Older participants and at more advanced pubertal stages were more frequently socially isolated (β = 0.04 to 0.07, CI = [0.002, 0.13], p < 0.05). Preferring solitude was associated with higher internalizing behaviors (β = 0.45, CI = [0.42, 0.49], p < 0.01), externalizing behaviors (β = 0.23, CI = [0.20, 0.27], p < 0.01), depression (β = 0.39, CI = [0.35, 0.42], p < 0.01), and anxiety (β = 0.29, CI = [0.26, 0.33], p < 0.01). Similarly, being socially withdrawn was associated with higher internalizing behavior (β = 0.41, CI = [0.37, 0.45], p < 0.01), externalizing behaviors (β = 0.25, CI = [0.21, 0.29], p < 0.01), depression (β = 0.45, CI = [0.42, 0.49], p < 0.01), and anxiety (β = 0.35, CI = [0.31, 0.39], p < 0.01).

Parental mental health issues were positively associated with youth preferring solitude and social withdrawal (β = 0.14, CI = [0.10, 0.19], p < 0.01, and β = 0.13, CI = [0.09, 0.17], p< 0.01 ), respectively). In contrast, being energized by large crowds of people was negatively associated with preferring solitude (β = -0.23, CI = [-0.27, -0.19], p < 0.01). When participants who often preferred solitude were excluded (due to their small sample size (n = 48)), and only participants who sometimes preferred solitude were compared to those who did not, additional positive associations were identified with being shy (β = 0.45, CI = [0.34, 0.55], p < 0.01), and not liking to meet new people (β = 0.58, CI = [0.46, 0.70], p < 0.01). Social withdrawal was also associated with being shy (β = 0.21, CI = [0.16, 0.25], p < 0.01), not liking to meet new people (β = 0.21, CI = [0.17, 0.25], p < 0.01), and feeling shy about meeting new people (β = 0.19, CI = [0.15, 0.23], p < 0.01).

### 3.2 Associations between social isolation and resting-state network properties

#### 3.2.1 Whole-brain topology

In the full cohort, there were no consistent associations across both fMRI runs. However, within-group comparisons of those who preferred solitude identified negative correlations between preference for solitude and connectome efficiency and global clustering (β = -0.03 to -0.02, CI = [-0.05, -0.01], p ≤ 0.04). Additional associations were estimated based on only the larger sample with the best-quality fMRI run. In the full cohort, preferring solitude was negatively associated with global efficiency (β = -0.07, CI = [-0.11, -0.03], p = 0.01), and similarly for pairwise group comparisons. Also, negative associations between preferring solitude and topological robustness, and similarly for stability, were estimated in group comparisons, and positive associations with modularity and segregation. There were no significant associations between brain-wide topological properties and being withdrawn. Statistics of models based only on the best run are summarized in Table S2.

#### 3.2.2 Network topology

Preferring solitude was associated with higher segregation (more connected local communities) of the right dorsal attention, temporoparietal, and social networks (β = 0.07 to 0.10, CI = [0.02, 0.14], p < 0.04). Within-group comparisons identified associations between preference for solitude and higher segregation of the right social network (β = 0.01, CI = [0.003, 0.01], p = 0.04) but lower global efficiency of the left cerebellum (β = -0.09, CI = [-0.13, -0.05], p < 0.01). In addition, social withdrawal was associated with lower topological robustness and stability (β = -0.78 to -0.67, CI = [-1.34, -0.19], p < 0.03) and higher fragility (β = 0.04, CI = [0.01, 0.07], p < 0.05) of the left cerebellum. Model statistics are summarized in Table 3.

**Table 3.** Statistics of mixed-effect models testing associations between preferring solitude, social withdrawal, and topological properties of individual networks. All reported p-values have been adjusted for the False Discovery Rate. Regression coefficients in models comparing pairs of groups were not standardized. *-: Nonsignificant; CI: confidence interval.

| Would rather be alone than with others |  |  |  |  |  |  |  |  |
| --- | --- | --- | --- | --- | --- | --- | --- | --- |
| Property | Network | Sample comparison | Beta | 95th % CI* | P-value | Beta | 95th % CI | P-value |
|  |  |  | Left hemisphere |  |  | Right hemisphere |  |  |
| Segregation | Dorsal attention | Entire sample | -* |  |  | 0.066 | [0.022, 0.110] | 0.034 |
|  | Social | Entire sample | - |  |  | 0.085 | [0.042, 0.128] | 0.001 |
|  |  | Somewhat vs Very True | - |  |  | 0.008 | [0.003, 0.013] | 0.037 |
|  | Temporoparietal | Entire sample | - |  |  | 0.095 | [0.051, 0.139] | <0.001 |
| Global efficiency | Cerebellum | Somewhat vs Very True | -0.089 | [-0.133, -0.045] | <0.001 | - |  |  |
| Withdrawn, does not get involved with others |  |  |  |  |  |  |  |  |
| Property | Network | Sample comparison | Beta | 95th % CI | P-value | Beta | 95th % CI | P-value |
|  |  |  | Left hemisphere |  |  | Right hemisphere |  |  |
| Topological robustness | Cerebellum | Somewhat vs Very True | -0.781 | [-1.342, -0.220] | 0.021 | - |  |  |
| Topological stability | Cerebellum | Somewhat vs Very True | -0.670 | [-1.151, -0.189] | 0.021 | - |  |  |
| Fragility | Cerebellum | Somewhat vs Very True | 0.039 | [0.005, 0.073] | 0.047 | - |  |  |

Additional associations were identified in analyses of only the best-quality fMRI. Preferring solitude was associated with higher segregation, modularity and/or fragility of multiple networks, including the basal ganglia, dorsal attention, prerontal, reward, salience, somatomotor, social and temporoparietal networks (β = 0.06 - 0.10, CI = [0.01, 0.14], p < 0.05). It was also associated with lower connectivity, topological robustness, stability and efficiency of most of the same networks, as well as the frontoparietal control network (β = -0.06 to -0.05, CI = [-0.10, -0.01], p < 0.05). Being withdrawn was also associated with higher segregation of the right temporoparietal network (β = -0.08, CI = [-0.03, -0.12], p < 0.01). Across networks, models on associations between preference for solitude and topological properties (including within-network median connectivity, fragility and global clustering) had good predictive power (CV[RMSE] ≤ 0.20). Similar results were obtained across pairwise group comparisons of those who strongly preferred solitude vs those who did not, within-group comparisons of those who preferred it, and those who somewhat preferred solitude vs those who did not, and similarly for social withdrawal. Detailed results and model statistics are provided in Table S3.

Finally, there were no consistent associations between social isolation and node properties across both fMRI runs. For the best-quality run, preferring solitude was associated with lower connectedness (degree) in nodes of the left dorsal attention network (β = -0.08 to -0.06, CI = [-0.12, -0.02], p <= 0.04). There were no other significant associations.

### 3.3 Associations between social withdrawal and structural properties

Preferring solitude was associated with lower thickness of the left superior temporal gyrus, right caudal anterior cingulate gyrus, right insula, and the left cuneus and pars opercularis (β = -0.09 to -0.05, CI = [-0.14, -0.01], p < 0.05), and lower white matter intensity but higher volume of the right parahippocampal gyrus (β = -0.02, CI = [-0.04, -0.002], p < 0.05, and β = 0.06, CI = [0.01, 0.10], p < 0.05 respectively). Social withdrawal was associated with higher white matter intensity in the right isthmus of the cingulate gyrus (β = 0.03, CI = [0.01, 0.05], p = 0.04). These associations were based on models with good predictive power (CV[RMSE] ≤0.20).

Comparisons of those who strongly preferred solitude to those who did not, identified associations between preference for solitude and lower thickness of the left cuneus, and left superior temporal gyrus, right caudal anterior cingulate gyrus, and right insula (β = -0.04 to -0.02, CI = [-0.07, -0.003], p < 0.05), lower volume of the left cuneus and right fusiform gyrus (β = -264.31 to -109.54, CI = [-474.51, -11.40], p = 0.04), and lower white matter intensity of the right parahippocampal gyrus (β = -0.23, CI = [-0.42, -0.05], p = 0.04). Social withdrawal was associated with higher volume of the right pallidum (β = 81.06, CI = [12.50, 149.62], p = 0.02), lower thickness of the right cuneus, right insula , and lingual gyrus (β = -0.06, CI = [-0.09, -0.01], p ≤ 0.04), and lower volume of the right cuneus (β = -333.56, CI = [-529.69, -137.44], p < 0.01) and right lingual gyrus (β = -492.48, CI = [-848.47, -136.49], p = 0.02).

In within group comparisons, preference for solitude was associated with lower white matter intensity in the right inferior temporal gyrus (β = -0.39, CI = [-0.68, -0.10], p = 0.03), and social withdrawal was associated with higher volume of left caudate nucleus, right pallidum and the putamen (β = 231.75 -369.01, CI = [2.68, 668.10], p < 0.05), lower thickness of the right cuneus (β = -0.15, CI = [-0.23, -0.06], p < 0.01) and right lingual gyrus (β = -0.09, CI = [-0.17, -0.01], p = 0.04), and lower volume of the right cuneus (β = -723.26, CI = [-1185.41, -261.12], p < 0.01).

Comparisons of those who somewhat preferred solitude versus those who did not, identified associations between preference for solitude and higher volume of the left amygdala, and right entorhinal and parahippocampal gyri (β = 25.72 -61.62, CI = [0.13, 106.57 ], p < 0.05), and lower thickness of the right pars opercularis and right caudal anterior cinguate gyrus (β = -0.03, CI = [-0.06, -0.003], p < 0.05). Similar comparisons of those who were somewhat socially withdrawn to those who were not, identified a negative association between social withdrawal and volume in of the right caudal middle frontal gyrus (β = -319.88, CI = [-579.79, -59.97], p < 0.05). Model statistics are summarized in Table 4. Negative associations between preference for solitude and cortical thickness, which were more consistent across group comparisons (especially in the caudal anterior cingulate cortex) are shown in Figure 1.

**Figure 1.**
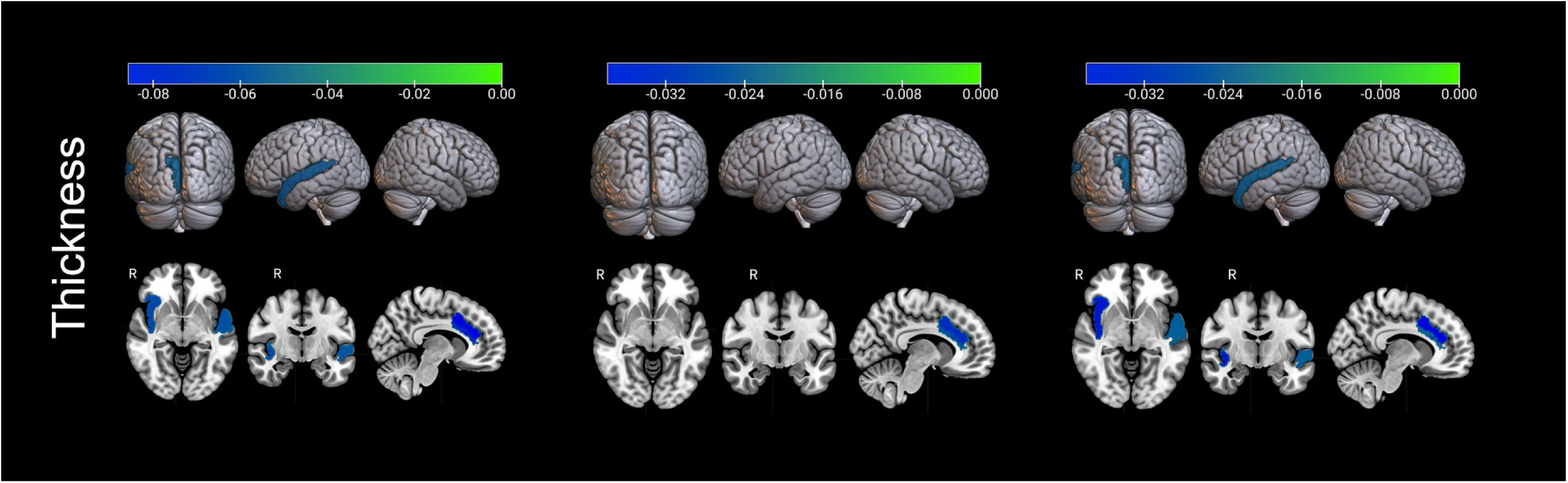
Associations between preferring solitude and cortical thickness. The color map represents the regression coefficients corresponding to cortical thickness.

**Table 4.**
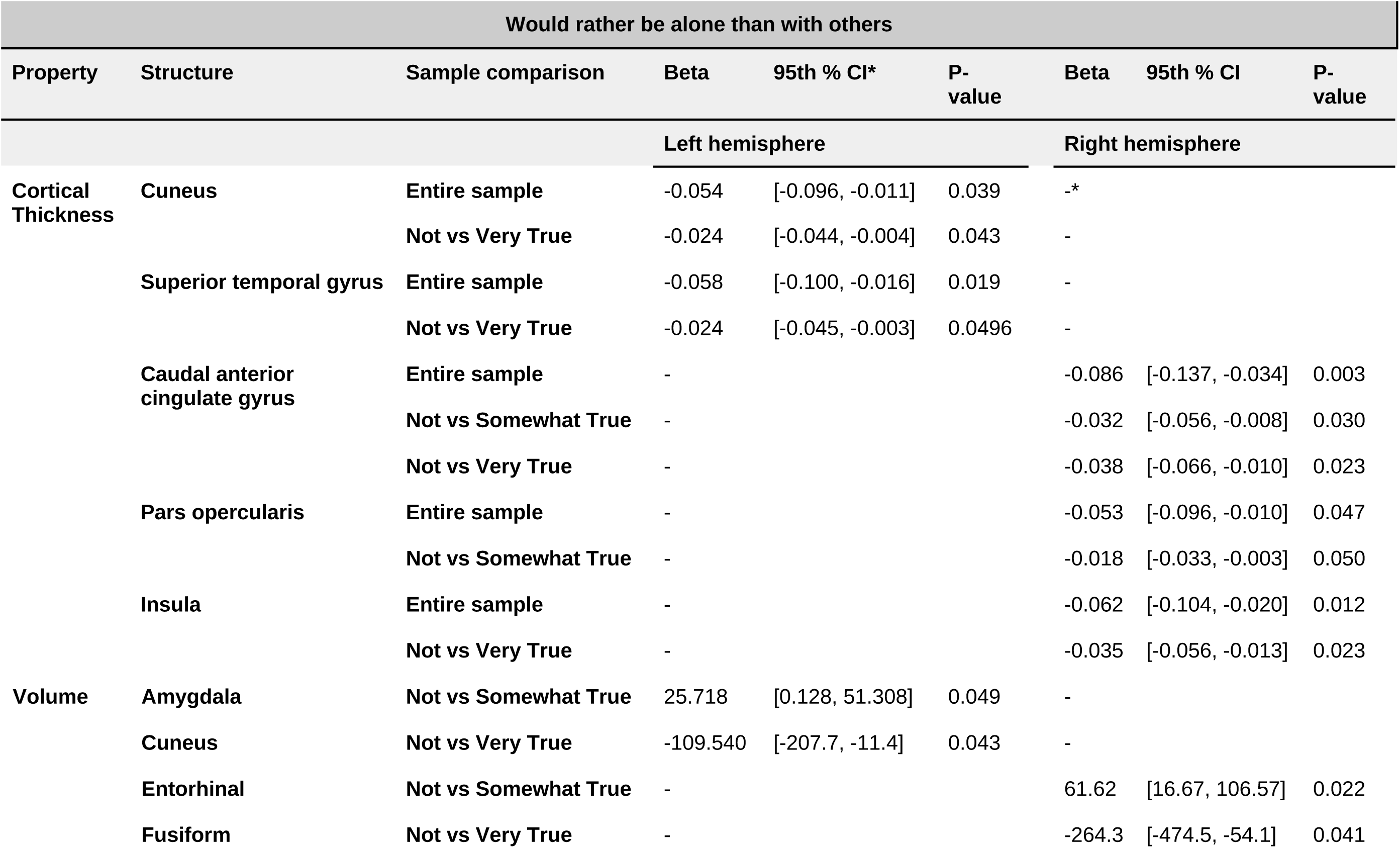
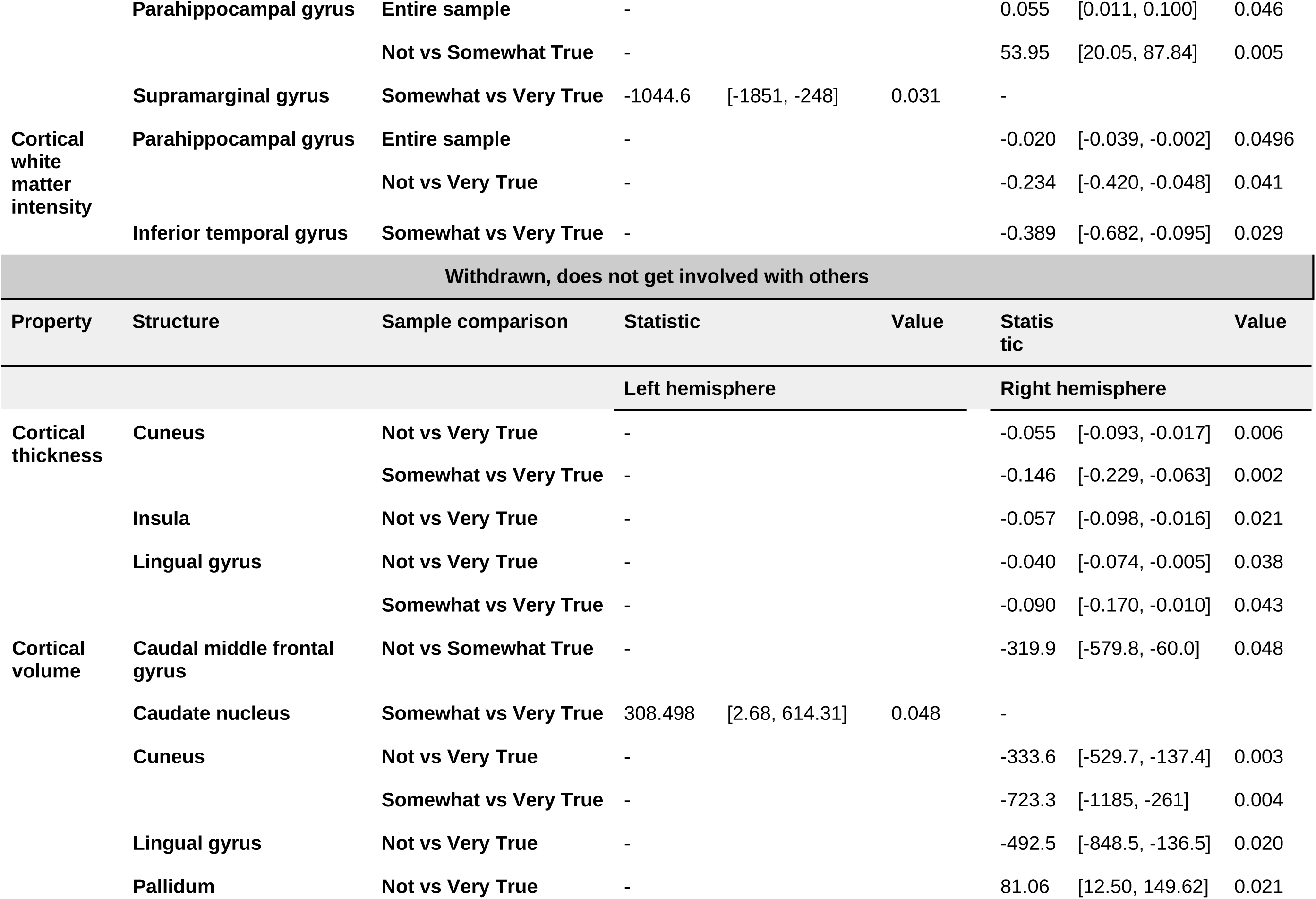
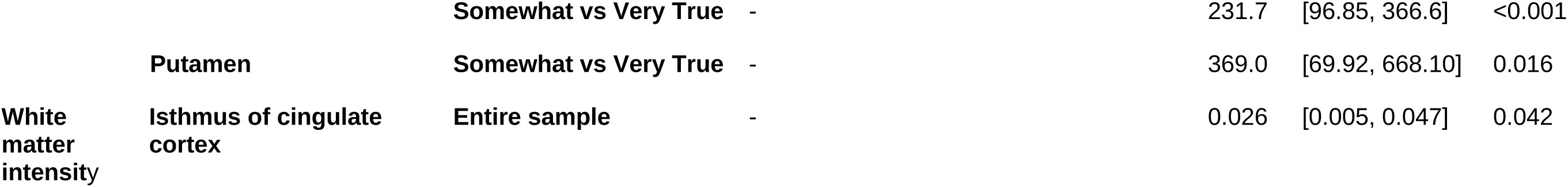
Statistics of mixed-effect models testing associations between preferring solitude and morphological properties. All reported p-values have been adjusted for the False Discovery Rate. Regression coefficients in models comparing only 2 response options were not standardized. *-: Nonsignificant *CI: confidence interval.

### 3.4 Mappings between functional and structural correlates of preference for solitude

Segregation of the right dorsal attention network was associated with morphometric properties of the right pars opercularis (β = 0.06, CI = [0.02, 0.11], p < 0.02) and the right inferior temporal gyrus (β = -0.06, CI = [-0.12, 0.01], p < 0.04), which both overlap with this network and were negatively associated with preference for solitude. Similarly, segregation of the right social network was associated with morphometric properties of the overlapping right insula (β = 0.08, CI = [0.02, 0.13], p < 0.03), and preference for solitude was associated with higher segregation of the network and lower thickness of the region.

## 4. DISCUSSION

In a large cohort of almost 3,000 adolescents, we have systematically investigated the neural correlates of preference for solitude and social withdrawal, especially their impacts on hallmark structural and topological characteristics of large scale networks that play critical roles in cognitive (including social) function. Social isolation in adolescence, a sensitive period of brain and social development but also vulnerable period for mental health, may lead to brain circuit miswiring, and increased risk of cognitive delays and deficits, and maladaptive behaviors across the lifespan. In this study’s cohort, about 18% of youth sometimes or often preferred solitude, and 6% were socially withdrawn. Older youth whose parents had mental health issues were more frequently preferred solitude and/or were socially withdrawn. These findings are in agreement with prior studies that have shown that parental behaviors and their mental health issues are significant risk factors for youth loneliness, social withdrawal and isolation (Matthews et al, 2015; Almeida et al, 2021), as early as infancy (Mantymaa et al, 2008).

As their social world expands beyond the family, and changes as a function of age, adolescents may become increasingly exposed to additional social stressors, such as peer rejection and victimization, which may, in turn, lead to social withdrawal and isolation. In this cohort, almost 90% had a regular group of friends, only ∼10% had experienced discrimination (or any form), and ∼20% had been bullied. Neither overall discrimination nor bullying were statistically associated with an increased likelihood of social isolation. However, unmeasured (by the ABCD) granular aspects of peer relationships and their quality and other latent factors in the youth environment that change with age could contribute to social isolation. Preference for solitude and social withdrawal was also associated with youth depression, anxiety, and internalizing and externalizing behaviors, in agreement with prior work that has identified extensive links between social isolation and mental health issues in youth (Rubin et al, 2009; Rubin and Lollis, 2015; Orben et al, 2020; Christiansen et al, 2021).

Preferring solitude and social withdrawal had overlapping structural but distinct topological brain correlates. Both preference for solitude and social withdrawal were associated with lower thickness of the cuneus and insula, while preference for solitude was also associated with lower thickness of the superior temporal and caudal anterior cingulate gyri and the pars opercularis, and social withdrawal with lower thickness of the lingual gyrus. The cuneus is a structural brain hub (Oldham and Fornito, 2019), i.e., a brain region where multimodal information is integrated, and has been implicated both in emotional processing and social function (Riedel et al 2018; Eslinger et al, 2021), but also loneliness (Lam et al, 2021; Wong et al, 2022). The insular cortex is another structural hub involved in information integration and supports interoception, but is also involved in (social) emotional processing and maladaptive social behaviors (Lamm and Singer, 2010; Terasawa et al, 2013; Emmerling et al, 2016; Gogolla, 2017). The superior temporal gyrus is considered a ‘hub’ of the social brain and is extensively involved in social perception and processing (Blakemore, 2008; Pelphrey and Carter, 2008; Lahnakoski et et al, 2012; Beauchamp, 2015). Decreased thickness of the cuneus, insula, and superior temporal gyrus has been associated with cognitive deficits but also mental health issues, not only in adults but also adolescents (von Tol et al, 2014; Pan, 2015; Sheffield et al, 2021). The anterior cingulate cortex has distributed connections with frontal and limbic regions and plays a ubiquitous role in cognitive function, including decision-making, cognitive control, and emotion and reward processing and regulation, and is also involved in social cognition (Devinsky et al, 1995; Vogt, 2005; Apps et al, 2016). The caudal part of the anterior cingulate cortex is involved in cognitive control, and decreased thickness of this region has been associated with depression, and developmental disorders affecting social function, such as Autism Spectrum Disorders (ASD) (Laidi et al, 2019; Mertse et al, 2022). Finally, the pars opercularis and lingual gyrus are both involved in speech and language processing (Palejwala et al, 2021; Zaccarella et al, 2015), and morphological anomalies in these regions have been associated both with linguistic deficits but more broadly impaired social communication, and mental health issues, including depression and internalizing symptoms (Jensen et al, 2015; Rosada et al, 2023).

Preference for solitude and social withdrawal were also associated with other structural differences in some of the same areas, including lower volume of the cuneus and the lingual gurys, but also higher volume of the entorhinal and parahippocampal gyri, amygdala and basal ganglia structures, including the caudate, pallidum and putamen. These findings are in agreement with those of prior studies that have associated social isolation and loneliness with higher amygdala volume (Xiong et al, 2023), potentially as a result of higher emotional sensitivity, but also distress in social settings (Lam et al, 2021; Vitale et al, 2022) and mental health issues (Espinoza et al, 2020; Suor et al, 2024). Specifically in children and adolescents, it has been associated with anxiety, negative affect and behavioral issues (Merz et al, 2018; Liu et al, 2024; Pereira et al, 2024). In addition, higher putamen volume has been associated with disorders affecting social function, such as ASD (Sato et al, 2014), and higher caudate volume with behavioral disregulation, impulsivity, and risk for developmental disorders such as ASD (Voelbel et al, 2006). Together these findings suggest that social isolation is associated with extensive morphological alterations in the adolescent brain in regions supporting social function but that have also been implicated in mental health disorders and social dysfunction, and developmental disorders impacting social communication. In addition, cortical thinning and increased structural volume are hallmark characteristics of brain development. It is possible that social isolation may be associated with accelerated neural maturation in selective regions, which, in turn, increases the risk for mental health and behavioral problems. A recent study on adolescents during the COVID-19 pandemic reported accelerated brain maturation partly as a result of social isolation due to lockdowns and social distancing (Corrigan et al, 2024).

Preference for solitude was also associated with higher topological segregation (modularity) of the dorsal attention and temporoparietal networks, and an overlapping distributed network representing the social brain (Blakemore, 2008). Increased structural and functional network modularity is also a fundamental characteristic of brain and cognitive development (Baum et al, 2017; Tooley et al, 2022), however in socially isolated youth it may reflect aberrantly accelerated neural maturation. Higher modularity has also been associated with mental health issues (Gao et al, 2023). Furthermore, social withdrawal was associated with topological changes in the cerebellum, including lower robustness and efficiency and higher fragility. A number of studies have shown that the cerebellum, which has extensive cortical and subcortical connections with brain regions supporting social function, plays an important role in social development, perception and prediction, and emotional processing) (Van Overwalle, 2020, 2024; Turrini and Avenanti, 2024). Abnormal structural changes in the cerebellum have been associated with developmental and mental health disorders, including those directing affecting social function (Fett et al, 2014; Jack and Morris, 2014; Phillips et al, 2015; Zhu and Qiu, 2022; Olivito et al, 2023; Wang et al, 2023). Together, these findings suggest that social isolation during adolescence may have profound detrimental impacts on large-scale networks and their constituent structures that play ubiquitous roles in cognitive, and specifically social function, and are abnormally modulated by mental health disorders.

Despite its strengths, including the large sample that captures the heterogeneity of adolescent brain development, comprehensive assessment of topological properties of resting-state networks, concurrent investigation of morphological and functional brain characteristics, and examination of environmental correlates of youth social isolation, this study also had some limitations. First, assessments of social withdrawal and preference for solitude were based on parent reports. When possible, youth surveys were analyzed, but across surveys measuring the youth environment and mental health/behavioral assessments, parent reports were in general more complete and reliable, or the only available reports (corresponding youth ones were not available).

Second, as is the case with any retrospective investigation, analyses were limited by the experimental decisions made by the ABCD investigators. Thus, more granular assessments of the youth social environment and related behaviors were not available. Nevertheless, the ABCD is the only investigation that extensively samples the youth social world, and relationships with parents, family, teachers, and peers. The present study leveraged data on these relationships in order to examine their associations with social isolation.

This study makes a significant scientific contribution and provides novel insights into the detrimental effects of social isolation on the developing adolescent brain and its maturating, and vulnerable to miswiring, neural circuitry. It has identified widespread morphological and topological brain alterations in youth who prefer to be alone and/or are socially withdrawn, several of which have not been previously related directly to social isolation, but to disorders affecting social communication. It has also associated preference for solitude and social withdrawal with parental mental health but also common youth mental health issues. Adolescence is a sensitive period for mental health. Many identified brain alterations were in regions that support social function but have also been implicated in mental health disorders. In addition, lower cortical thickness and higher subcortical volume and increased modularity of several large-scale networks also suggest potential accelerated maturation of the socially isolated brain, which has also been associated with mental health issues. The ABCD follows youth longitudinally, thus future investigations could specifically examine the hypothesis of accelerated neural maturation and track the emergence of mental health issues as a function of age in socially withdrawn youth.

### Data Availability Statement

The data underlying this article are available in National Institute of Mental Health Data Archive (NDA). https://nda.nih.gov/.

## Acknowledgments

This study was supported by the National Science Foundation, grant numbers 2207733, 2116707, 1940096.

